# Alkaline phosphatase activity supports heterotrophic carbon acquisition in a coastal time series site and a representative marine bacterium

**DOI:** 10.64898/2026.03.24.713987

**Authors:** Eesha Sachdev, Jamee C. Adams, Kaycie B. Lanpher, Shannon Perry, Carina Tostado, Jeff S. Bowman, Ellery D. Ingall, Julia M. Diaz

## Abstract

Phosphorus is a vital nutrient required for the functioning of living organisms. In aquatic environments, dissolved inorganic phosphate is considered its most bioavailable form. However, phosphate can be scarce, which has the potential to limit microbial metabolism and ecosystem functioning. To overcome phosphate scarcity, microbes produce alkaline phosphatase (AP) to access dissolved organic phosphorus (DOP). Here, we conducted a year-long study of alkaline phosphatase activity (APA) at the Ellen Browning Scripps Memorial Pier, a nutrient-rich coastal site. APA was observed throughout the year despite phosphate-replete conditions, suggesting that the role of APs in microbial nutrition is not completely understood. We tested the hypothesis that APA may promote acquisition of organic carbon liberated from DOP hydrolysis by growing the heterotrophic marine bacterium *Ruegeria pomeroyi* on three DOP compounds as sole carbon sources and assessing APA. Controlling for carbon concentration, all DOP sources supported growth, but at lower levels than glucose, with the highest growth observed on glucose-6-phosphate (G6P), followed by adenosine monophosphate (AMP) and adenosine triphosphate (ATP). Moreover, cell-specific APA was significantly enhanced in carbon-deplete conditions and during growth on G6P, relative to cultures grown on replete glucose or nucleotides. These findings suggest alkaline phosphatases (APs) are part of a generic carbon stress response and likely play a role in acquiring certain forms of organic carbon by *R. pomeroyi*, with implications for other taxa. Overall, this study helps advance the current state of knowledge regarding microbial phosphorus cycling and carbon utilization in aquatic environments.

## Introduction

Phosphorus is an essential nutrient that plays a crucial role in the functioning of all living organisms through its presence in numerous biological processes and structures. It is vital for the expression of genetic information in biomolecules such as nucleic acids, DNA, and RNA. Additionally, phosphorus is involved in energy storage and transfer through adenosine triphosphate (ATP) and serves as a fundamental component of structural molecules like phospholipids (Karl, 2014).

In aquatic systems, microbial communities control the fate of phosphorus, carbon, and other bioactive elements. Phosphorus exists in both inorganic and organic forms. Dissolved inorganic phosphate is considered the most bioavailable form of phosphorus and is readily utilized by microbes (Mahaffey et al. 2014) compared to larger and more complex phosphorus-bearing organic molecules (Thomson et al. 2019). However, the distribution of phosphate in the ocean is not uniform, leading to regions of phosphorus scarcity compared to more nutrient-rich coastal and upwelling zones. This scarcity of phosphate can affect microbial metabolism, growth, and essential ecosystem functions (Karl, 2014).

Dissolved organic phosphorus (DOP) is a key nutritional resource, especially in low-phosphate regions (< 20 nM) such as the open ocean and subtropical gyres (Martiny et al. 2019). Microbes access DOP using the enzyme alkaline phosphatase (AP) (Duhamel et al. 2021; Su et al. 2023). AP activity (APA) is often used as an indicator of phosphorus nutritional status and interpreted to signal phosphate deficiency in a system (Duhamel et al. 2021; Hoppe, 2003; Su et al. 2023) because APA is thought to be inhibited by phosphate and upregulated when phosphate drops below critical thresholds (∼30 nM) (Dyhrman and Ruttenberg, 2006; Mahaffey et al. 2014; Sebastián et al. 2004). Yet APA has been routinely observed in phosphate-replete systems, such as coastal regions and deep waters (Davis and Mahaffey, 2017; Hoppe and Ullrich, 1999; Karl and Björkman, 2015; Koike and Nagata, 1997; Labry et al. 2016), a phenomenon which is referred to as the “APA enigma” or APA paradox (Karl, 2014). One explanation for the APA paradox in deep waters involves the export of surface-derived APs in fast-sinking particles (Davis and Mahaffey, 2017; Dykens et al. 2025; Koike and Nagata, 1997; Nausch and Nausch, 2004). Alternatively, it has been proposed that APs may be involved in the acquisition of organic carbon, rather than phosphate, liberated from the hydrolysis of bioavailable DOP (Hoppe and Ullrich, 1999; Lidbury et al. 2022; Nicholson et al. 2006; Saavedra et al. 2025).

AP targets DOP by hydrolyzing the phosphate group from the organic molecule (Dyhrman and Ruttenberg, 2006) (Fig. 1). APs are commonly known for hydrolyzing P-ester bonds (P-O-C), which can occur as monoesters (P-O-C) and diesters (C-O-P-O-C) (Hoppe and Ullrich, 1999; Hoppe, 2003; Karl and Björkman, 2015; Mahaffey et al. 2014). P-esters represent approximately 80% of the marine DOP pool, with highly labile examples including adenosine monophosphate (AMP) and glucose-6-phosphate (G6P), which are both P-monoesters (Young and Ingall, 2010). APs can also hydrolyze P-anhydride bonds (P-O-P), which make up about 10% of the DOP pool (Adams et al. 2022; Duhamel et al. 2011; Young and Ingall, 2010). Adenosine triphosphate (ATP) is an example of a DOP source containing both P-ester and P-anhydride bonds.

**Figure 1:**
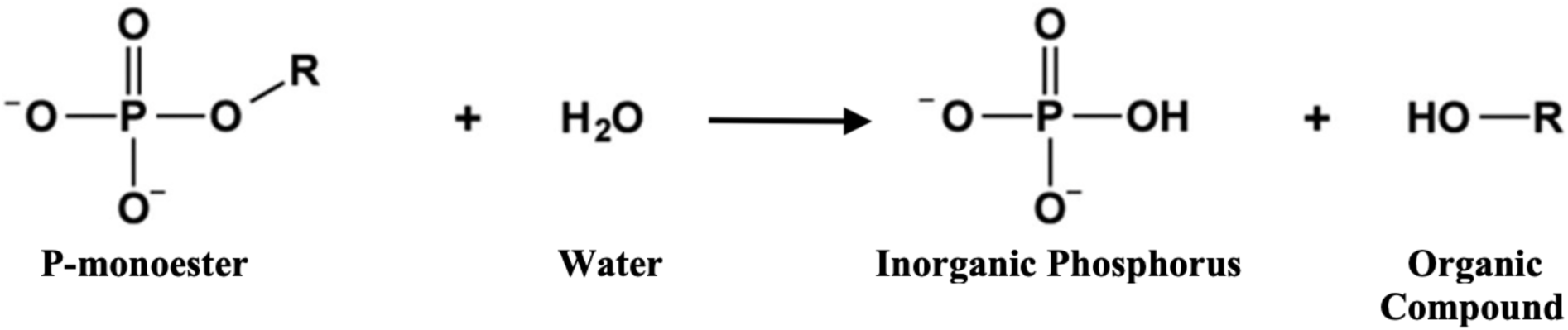
Chemical reaction performed by alkaline phosphatase. *R* represents the organic molecule bound to the phosphate group.

In this study, we explored the potential roles of alkaline phosphatase in phosphate-replete marine systems and investigated the relationship between DOP bioavailability and carbon acquisition. First, we conducted a year-long analysis of APA at the Ellen Browning Scripps Memorial Pier to determine whether, and to what degree, APA would be observed in this relatively high-nutrient coastal system. Next, we tested the model coastal marine heterotrophic bacterium *Ruegeria pomeroyi*, a member of the Roseobacter clade of Alphaproteobacteria, to determine whether it can use various DOP sources as a sole carbon source.

## Methods

### Field sampling

Seawater samples were collected from the Ellen Browning Scripps Memorial Pier (32°52′N, 117°15.4′W) every Monday and Thursday between 11:45 am to 12:00 pm local time, from October 2022 to October 2023. Depending on weather conditions, either the indoor sampling room with a hand crank or the outdoor crane was used for sample collection. All sampling containers were acid-cleaned in the lab prior to use. Immediately before sample collection, containers were rinsed three times with seawater. All samples were filtered (250 µm) to remove large particulate matter or debris before being transferred into 1-liter bottles for further processing. Seawater samples for soluble reactive phosphorus (SRP) and DOP were filtered using pre-combusted (450°C for 5 hours) GF75 filters (0.3 µm nominal pore size) and frozen in sterile 50 mL conical tubes at -20°C until analysis. Dissolved organic carbon (DOC) samples were collected following established protocols (Carlson et al. 2010; Knap et al. 1996). Briefly, seawater was filtered through a pre-combusted (450°C for 5 hours) and acid washed (10% HCl) GF/F filter (0.7 µm nominal pore size) into a combusted (450°C for 5 hours) glass vial, acidified with 50 µL of 4N HCl (final pH ∼3), and stored at room temperature until analysis. Chlorophyll-*a* samples were collected into opaque bottles, collected on GF/F filters (0.7 µm nominal pore size) in the dark, wrapped in foil to protect from light, and stored at -80°C until analysis. Unfiltered water for APA analysis was transported to the lab and promptly analyzed.

### Culture experiments

*Ruegeria pomeroyi* DSS-3 was grown in defined media modified from the recipe by Rivers et al (2016). To prepare media, 100 mL of filtered (0.2 µm) natural seawater collected from the Scripps Pier was autoclaved (121°C, 20 mins) in each of several acid washed 125 mL borosilicate flasks. Filter-sterilized (0.2 µm) nutrients and carbon sources were added in a laminar flow hood using aseptic techniques. Glucose, with a final concentration of 27 mM C, served as the carbon source for a C-replete control (10× glucose). The experimental DOP sources (Fig. 2), glucose-6-phosphate (G6P), adenosine monophosphate (AMP), and adenosine triphosphate (ATP), were prepared to a final concentration of 27 mM C. All DOP-containing media were supplemented with 2.7 mM of C as glucose (final) to support initial growth. A C-deplete control was prepared with a final concentration of 2.7 mM of C as glucose (1× glucose). All treatments had 250 µM of phosphate in the initial media.

**Figure 2:**
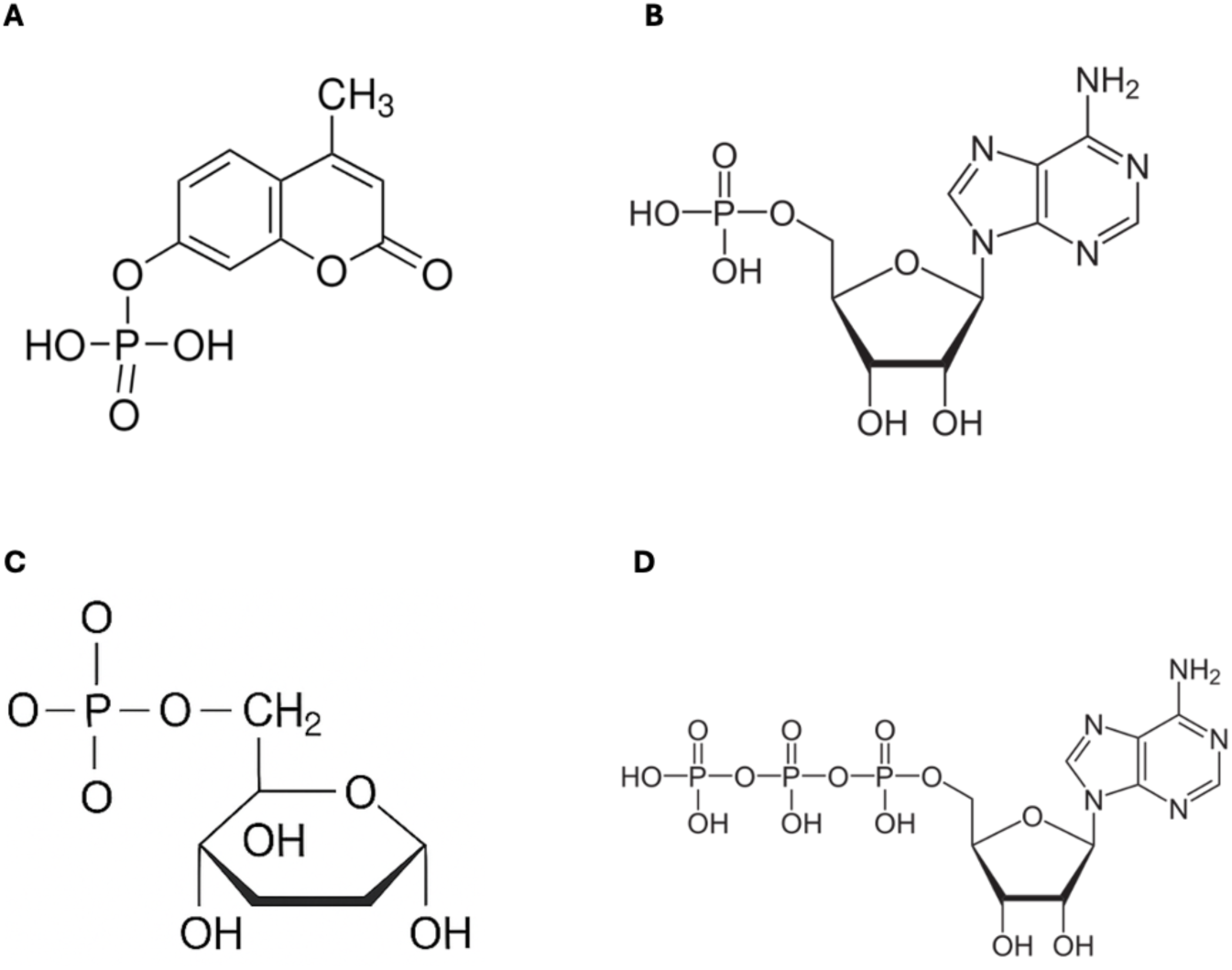
DOP sources used in this study. **(A)** 4-methylumbelliferyl phosphate (MUF-P) **(B)** Adenosine monophosphate (AMP) **(C)** Glucose-6-phosphate (G6P) **(D)** Adenosine triphosphate (ATP)

Each flask was inoculated with 50 µL of *R. pomeroyi* from a starter culture grown to exponential phase in C-replete media described above. Control and experimental groups were grown in triplicate in a Thermo Fisher MaxQ 6000 refrigerated shaker/incubator at 30°C, shaking at 150 RPM for 13 days.

Bacterial growth was monitored daily by measuring optical density at 600 nm using a SpectraMax M3 Multi-Mode Plate Reader (Molecular Devices). Optical density readings were used to calculate growth rates (d^-1^) as the slope of the log-linear period of growth (average R^2^ = 0.941). Growth rates were calculated from days 1-2 for C-replete (10× glucose), days 2-8 for C-deplete (1× glucose) and G6P cultures, days 3-8 for AMP cultures, and days 2-6 for ATP cultures. On each sampling day, a 1 mL subsample was fixed with 20 μL of 0.22 μm-filtered 25% glutaraldehyde (0.5% final concentration) for flow cytometry and stored at -80°C until analysis. Samples for SRP were filtered (0.22 μm) and stored at -20°C until analysis. APA was measured daily.

### Alkaline Phosphatase Activity (APA)

To assess APA, 4-methylumbelliferyl phosphate (MUF-P) was used as a model fluorogenic DOP substrate (Fig. 2A). Fluorescence of the stable reaction product 4-methylumbelliferone (MUF; EX: 359 nm, EM: 449 nm) was measured over time. Analyses were conducted in a 96-well opaque plate (black). For field measurements, each well received 190 µL of unfiltered sample and 10 µL of MUF-P stock solution or 10 µL of deionized water to achieve final MUF-P concentrations of 0, 0.05, 0.1, 0.5, 1, 2, 5, and 10 µM (Supplementary Fig. 1). For *R. pomeroyi* cultures, a single MUF-P concentration of 10 µM was used, based on initial experiments that this concentration was saturating (Supplementary Fig. 2). Each MUF-P concentration was conducted in triplicate and analyzed on a SpectraMax M3 Multi-Mode Plate Reader (Molecular Devices). For field samples, fluorescence was measured over a 12-hour period, with measurements taken every 5 minutes during the first hour and hourly thereafter. For *R. pomeroyi* cultures, measurements were taken every 15 minutes over a 1-hour period.

Analyses were calibrated using a standard series of MUF (0–10 µM) prepared in filtered and boiled seawater. MUF-P autohydrolysis was assessed using filtered and boiled natural seawater amended with the same MUF-P concentrations as above.

The data obtained from the plate reader were analyzed using Microsoft Excel. First, hydrolysis rates (nmol phosphate L^-1^ h^-1^), were determined by calculating the slope of fluorescence vs time in each well, applying the MUF calibration curve to each slope, and assuming a 1:1 molar ratio of MUF to phosphate. Any negative or outlier hydrolysis rates (v) were addressed as follows: If two replicates were negative and one was positive, all three points were excluded from further analysis. If two replicates were positive and one was negative, the negative point was removed. Observations for the 0 µM MUF-P concentration were never excluded, regardless of negative or outlier values.

For *R. pomeroyi*, APA (V_max_) is reported as the average hydrolysis rate (v) of biological triplicates calculated at a single saturating concentration of MUF-P (10 µM). For field samples, APA (V_max_) and the half-saturation constant (K_m_) were determined by using the Hanes-Woolf equation, a linearized version of the Michaelis-Menten equation: S:v = K_m_:V_max_ + S: V_max_, where S is the concentration of MUF-P and v is the hydrolysis rate. Linear regression analysis was performed using the LINEST function with the least squares method.

### Soluble Reactive Phosphorus (SRP)

SRP analysis was performed following the colorimetric protocol outlined by Strickland and Parsons (1972) and Johnson (1971). Frozen samples stored in 15 mL conical tubes were thawed and treated with an arsenate reagent to prevent any potential interference. Samples were analyzed on a SpectraMax M3 Multimode Plate Reader (Molecular Devices) by measuring absorbance at 880 nm. SRP analysis was performed in triplicate.

### Dissolved Inorganic Nitrogen (DIN)

DIN data at the Scripps Pier were obtained from the Southern California Coastal Ocean Observing System (SCCOOS) Harmful Algal Bloom Monitoring and Alert Program (HABMAP). Samples were collected and analyzed using standard oceanographic protocols described in Hatch et al. (2013).

### Dissolved Organic Phosphorus (DOP)

DOP analysis was carried out following the protocol outlined by Monaghan and Ruttenberg (1999) and Solórzano and Sharp (1980). Frozen samples stored in 15 mL conical tubes were thawed, acidified with HCl (∼pH 1) and stirred overnight to remove any possible phosphorus from the tube walls. Samples were then mixed with 0.17 M MgSO_4_, dried overnight in an oven (95°C), and baked in a muffle furnace (500°C, 4 hours). Finally, samples were acidified with 0.75M HCl (80°C, 20 min), diluted with Milli-Q water (17 mL), heated (80°C, 10 min) and analyzed as SRP on a SpectraMax M3 Multimode Plate Reader (Molecular Devices) to determine Total Dissolved Phosphorus (TDP). TDP concentrations were used to calculate DOP using the equation, DOP = TDP - SRP.

### Dissolved Organic Carbon (DOC)

DOC samples were analyzed using a DOC analyzer (Shimadzu), according to established protocols (Benner and Strom 1993; Grasshoff et al. 1999). Replicate DOC samples typically agreed to within ±7%.

### Chlorophyll-a

Chlorophyll-*a* analysis was based on the protocols established by Arar and Collins (1997) and Strickland and Parsons (1972). After thawing, filters were placed in 10 mL of 90% acetone and extracted overnight at 4°C in the dark. The next day, samples were centrifuged (1000 × g, 5 min, 23°C) and fluorescence (Ex: 672 nm, Em: 660 nm) was measured in the supernatant using a SpectraMax M3 Multi-Mode Plate Reader (Molecular Devices). To minimize light exposure, samples were analyzed under low-light conditions and covered with aluminum foil when not in use. A standard curve (0–1000 µg/L) of chlorophyll-*a* (Sigma C6144) in 90% acetone was used to calibrate the analysis.

### Flow cytometry

*R. pomeroyi* samples were thawed to room temperature, and 20 µL was pipetted into a clear, round-bottom 96-well plate. Samples were stained with SYBR Green according to manufacturer instructions (Invitrogen, REF S7563). Samples were then diluted 1:10 with filtered (0.22 µm) and autoclaved (121°C, 20 minutes) seawater. Blanks were prepared in triplicate using filtered and autoclaved seawater amended with 0.22 µm filtered 25% glutaraldehyde (0.5% final concentration). Analysis was performed using a Guava EasyCyte HT flow cytometer (Cytek Biosciences) using a low flow rate (0.24 µL s⁻¹) for 3 minutes or 1000 events. Populations were gated using diagnostic signals of forward scatter versus green fluorescence.

### Statistical analysis

All statistical analyses were performed in R (version 4.4.1). Potential correlations between environmental parameters measured at the Scripps Pier were evaluated using simple linear regression. The effect of carbon source on *R. pomeroyi* growth yields or growth rates was analyzed using Tukey’s honest significant difference (HSD).

## Results

### Scripps Pier Time Series

Dissolved inorganic P, or SRP, fluctuated between ∼40 and 700 nM throughout the time series, reaching a maximum on January 22 and a minimum on April 20 (Fig. 3A). DIN ranged from 0.22 to 6.93 µM, peaking around the same time as SRP on January 17 (6.93 µM) and reaching its lowest value on August 22 (0.22 µM) (Fig. 3B). Molar ratios of DIN:SRP were typically below the Redfield value (16), with two observed exceptions in July (Fig. 3C).

**Figure 3:**
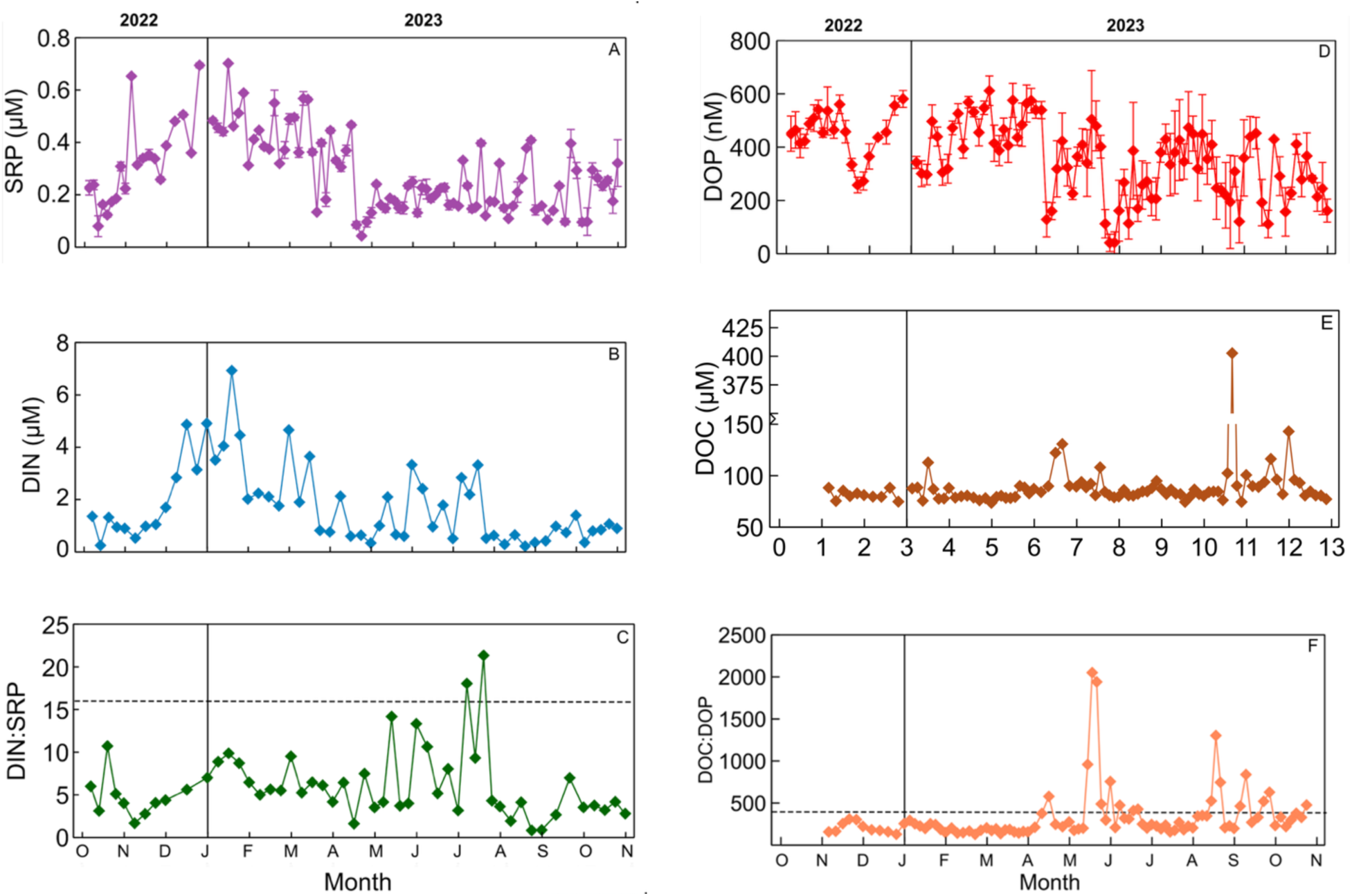
Dissolved carbon, nitrogen, and phosphorus at the Scripps Pier. **(A)** Soluble reactive phosphorus (SRP). Error bars represent the standard deviation of the mean of triplicate samples. **(B)** Dissolved inorganic nitrogen (DIN) **(C)** DIN:SRP molar ratio. Dashed line shows Redfield N:P = 16. **(D)** Dissolved organic phosphorus (DOP). Error bars represent the standard deviation of the mean of triplicate samples. **(E)** Dissolved organic carbon (DOC). Replicate DOC measurements typically agreed to within +/- 6.97%. **(F)** DOC:DOP molar ratio. Dashed line shows global DOC:DOP = 387 (Liang et al. 2023). Interpolation lines indicate linear interpolation.

DOP concentrations ranged from ∼40 to 600 nM, with the highest value on February 23 (612 nM) and the lowest on May 22 (41 nM) (Fig. 3D). DOP typically accounted for 56.5 ± 15.4% (± 1 sd) of the total dissolved P (Fig. 4F). In contrast, DOC remained relatively stable over the year, typically between 75 and 100 µM, with the highest concentration on August 24 (403 µM), and smaller peaks on October 2 (143 µM) and April 24 (131 µM) (Fig. 3E). The molar ratio of DOC:DOP peaked on May 22 (2,051) and reached a minimum on February 23 (128) (Fig. 3F). After removing the four highest outliers (May 22, May 25, Aug. 24, and Sep 14), the average DOC:DOP molar ratio was 278 ± 151 (± 1 sd).

**Figure 4:**
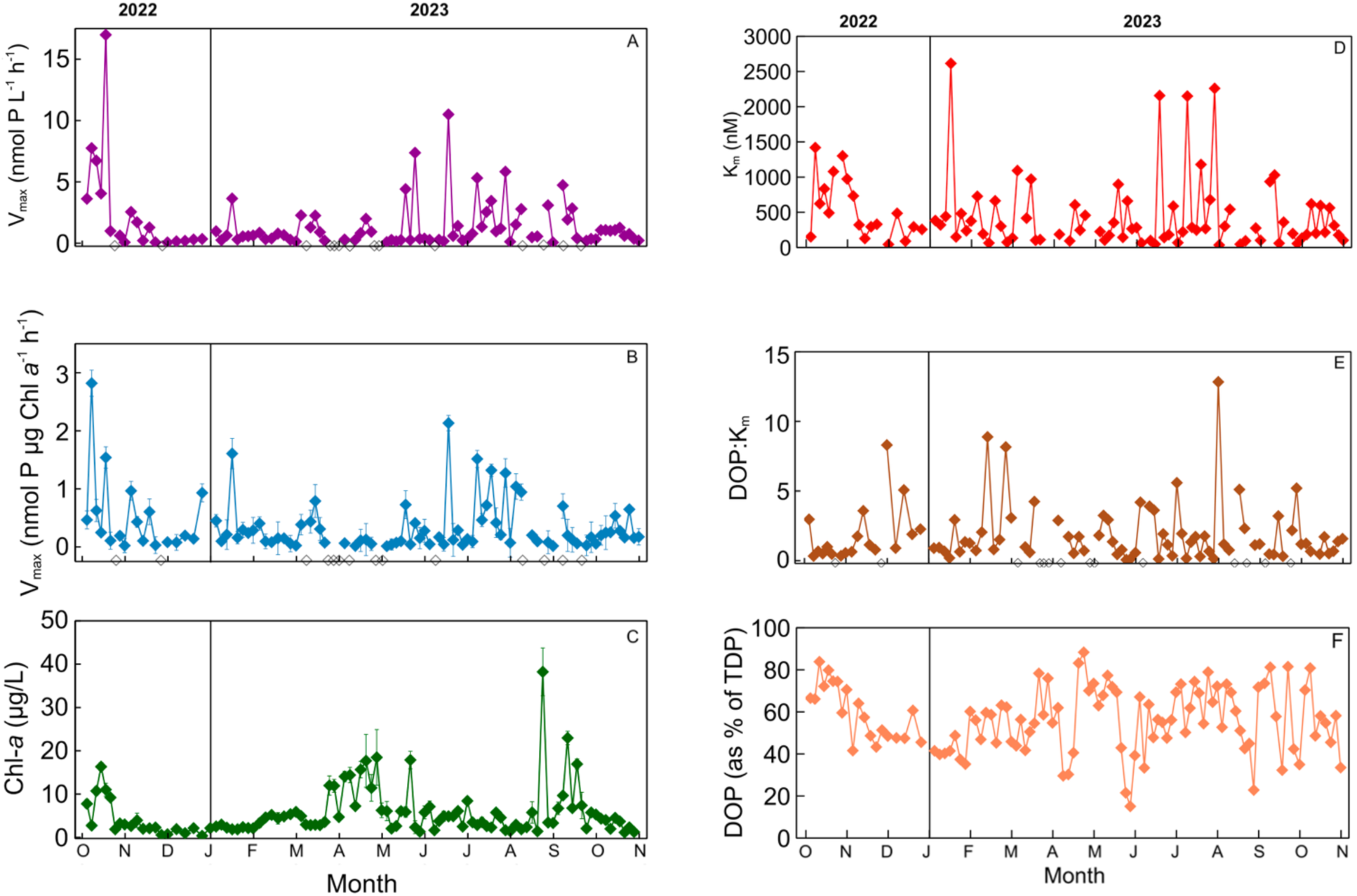
Alkaline phosphatase activity, chlorophyll, K_m_, K_m_:DOP and % DOP of TDP at the Scripps Pier. Maximum velocity of **(A)** volume-normalized and **(B)** chlorophyll-normalized alkaline phosphatase activity (V_max_). Open diamonds indicate data that were excluded from the results due to negative alkaline phosphatase activity. **(C)** Chlorophyll-*a*. **(D)** K_m_ **(E)** DOP:K_m_. **(F)** Percent DOP of TDP at the Scripps Pier. In all panels, error bars represent the standard deviation of the mean for triplicate samples. Interpolation lines indicate linear interpolation.

APA was observed throughout October 2022 to October 2023, with volume-normalized APA ranging from 0.06 to 16.99 nmol P L^-1^ h^-1^ (Fig. 4A). Chlorophyll-a varied over the year, peaking on August 24 (38.23 µg/L) and reaching a minimum on December 12 (0.37 µg/L) (Fig. 4C). APA (nmol P L^-1^ h^-1^) and chlorophyll were positively correlated (p=0.038), but chlorophyll explained only ∼5% of the variability in APA across the year (Supplementary Fig. 3).

APA data were normalized to chlorophyll-a concentration to account for variations in chlorophyll biomass (Fig. 4C). Chlorophyll-normalized APA ranged from 0.014 to 2.82 nmol P µg Chl-a⁻¹ h⁻¹, with the highest values on October 6, 2022 (2.82 nmol P µg Chl-a⁻¹ h⁻¹), October 17, 2022 (1.54), January 17 (1.61), June 15 (2.13), and throughout July and early August (1–1.5). APA generally remained below 0.500 nmol P µg Chl-a⁻¹ h⁻¹ for most of the sampling period. Fourteen days were excluded from the results due to negative V_max_ values (Fig. 4A, B).

K_m_ values ranged from 35 to 2615 nM (Fig. 4D), with a mean of 464 ± 508 nM (± 1 sd). The lowest observed value was on July 31 and the highest on January 17. DOP concentrations were higher than K_m_ for 54.5% of the observations over the time series (DOP: K_m_ >1) (Fig. 4E). V_max_ and K_m_ were significantly correlated (p = 2.72 × 10⁻¹⁴; Supplementary Fig. 4), but neither parameter showed significant relationships with SRP, DOP, or DOC (Supplementary Fig. 5).

### Culture Experiments

To determine whether bacteria can utilize DOP as a sole carbon source, *Ruegeria pomeroyi* was grown on three different DOP sources (27 mM C as AMP, ATP or G6P), each supplemented with 2.7 mM C as glucose. C-replete and C-deplete control cultures were grown on 10× glucose (27 mM C) or 1× glucose (2.7 mM C), respectively.

Cultures grown on 10× glucose reached a maximum optical density of 0.737 by day 3, while the cultures grown on 1× glucose only reached 0.086 (Fig. 5), confirming that growth was controlled by organic carbon availability. All DOP sources supported more growth than the 1× glucose control, including ATP (OD=0.214 on day 6), AMP (OD=0.314 on day 13) and G6P (OD=0.379 on day 13) (Fig. 5B).

**Figure 5:**
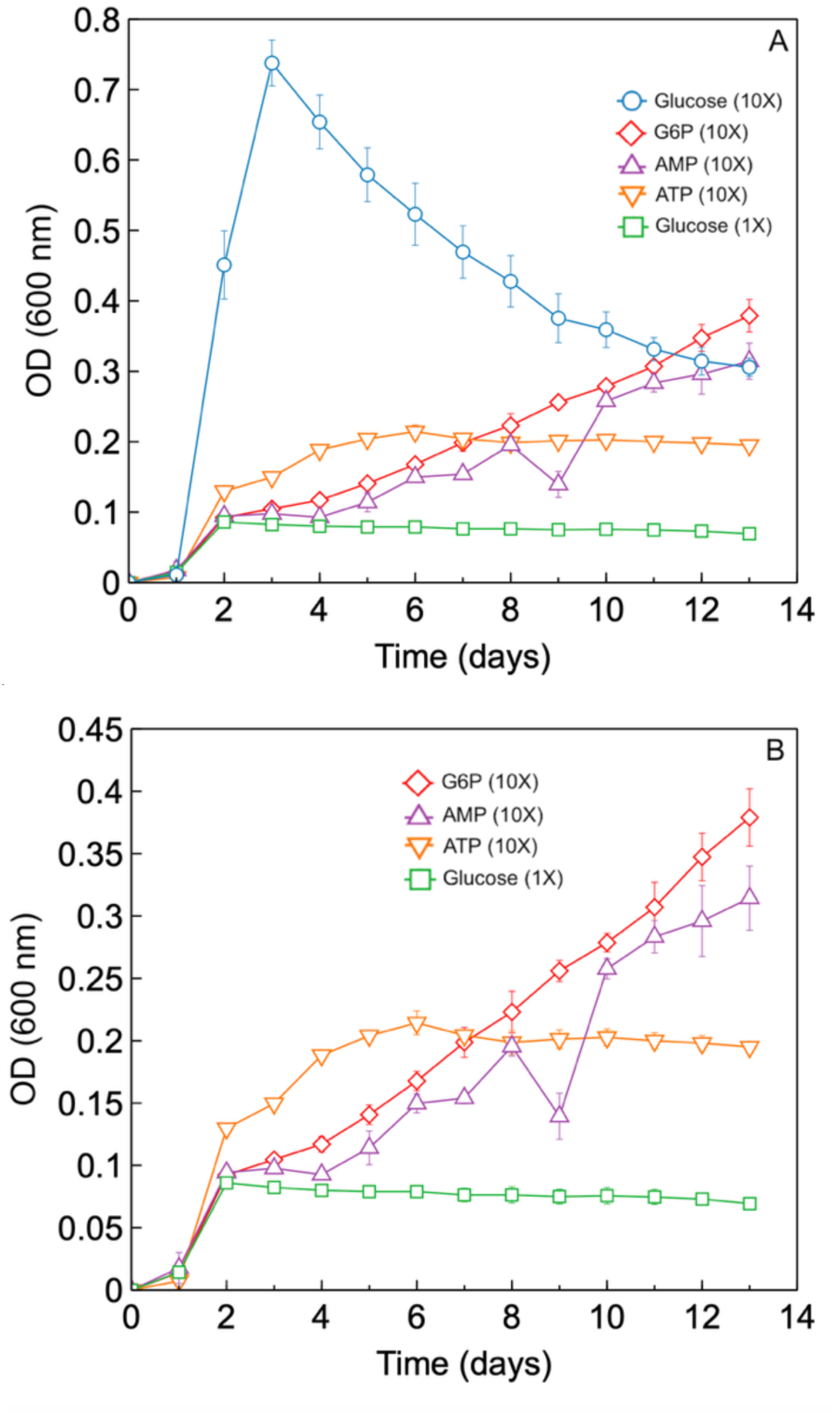
*R. pomeroyi* growth on DOP as a sole carbon source. **(A)** All treatments **(B)** With the 10× glucose group excluded for clarity. Error bars represent the standard deviation of the mean of biological triplicates. Interpolation lines indicate linear interpolation.

Final cell counts were largely consistent with optical density results. Final cell yields were highest for the 10× glucose cultures, reaching 2.94 × 10⁸ cells mL^-1^ (Fig. 6). Cell yields were 2.95 × 10⁶ cells mL^-1^ for the 1× glucose cultures, 1.93 × 10⁷ cells mL^-1^ for G6P, 1.64 × 10⁷ cells mL^-1^ for AMP, and 5.96 × 10⁶ cells mL^-1^ for ATP (Fig. 6). Tukey’s honest significant difference (HSD) test showed that G6P and AMP cultures did not differ significantly (p = 0.456), nor did ATP and 1× glucose cultures (p = 0.425). All remaining treatment comparisons were statistically significant (p < 0.01).

**Figure 6:**
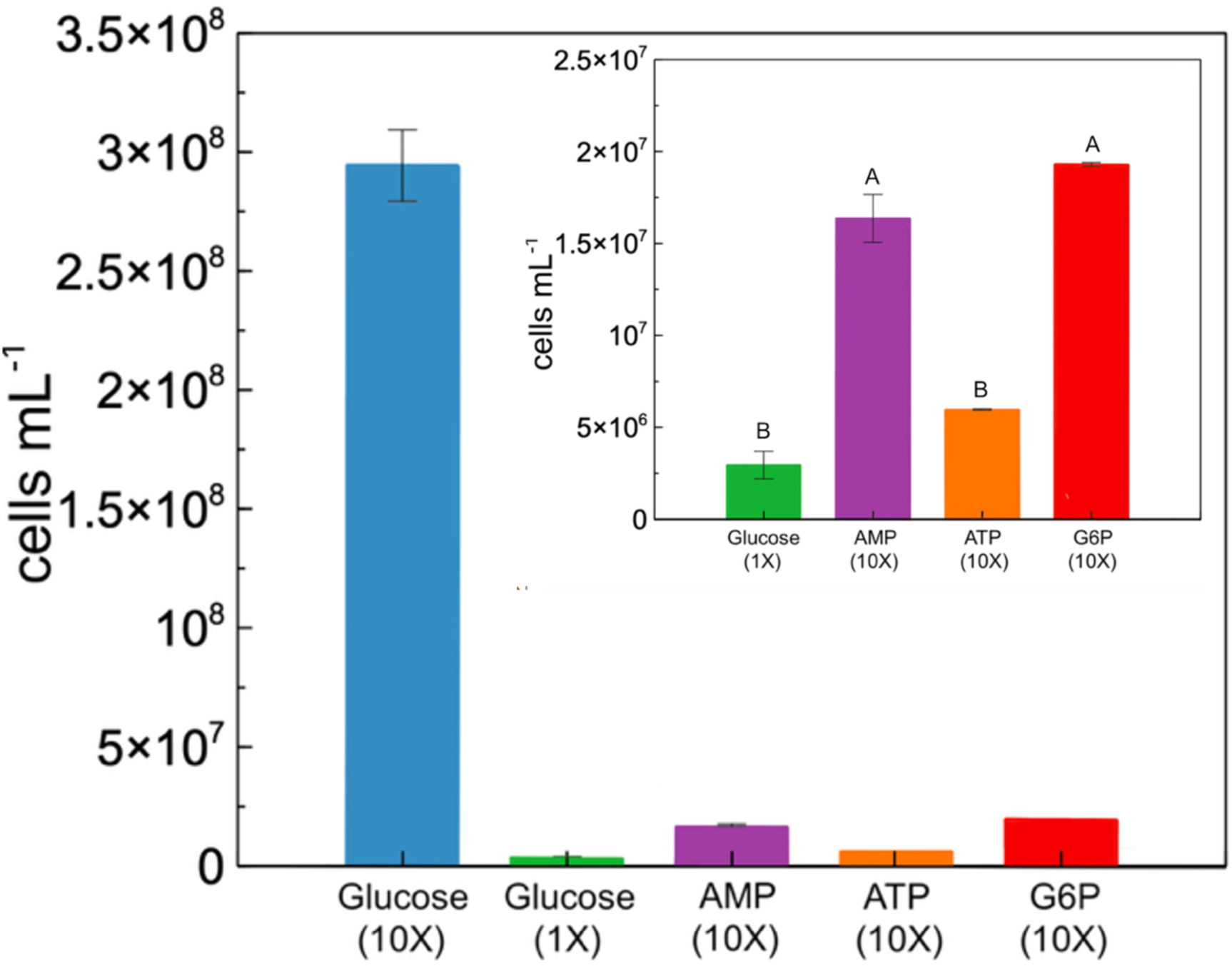
*R. pomeroyi* final cell yield on Day 13. Inset excludes the 10× glucose group for clarity. Groups lacking a shared letter are significantly different (p < 0.01) (Tukey HSD). Error bars represent the standard deviation of the mean for biological triplicates.

The growth rate for the 10× glucose cultures was 0.15 h⁻¹, while the 1× glucose cultures showed a net death rate of -0.0008 h⁻¹ (Fig. 7). Among the DOP groups, G6P (0.0064 h⁻¹), AMP (0.0062 h⁻¹), and ATP (0.054 h⁻¹) had similar growth rates, with no significant difference (p > 0.05) based on Tukey’s HSD test (Fig. 7).

**Figure 7:**
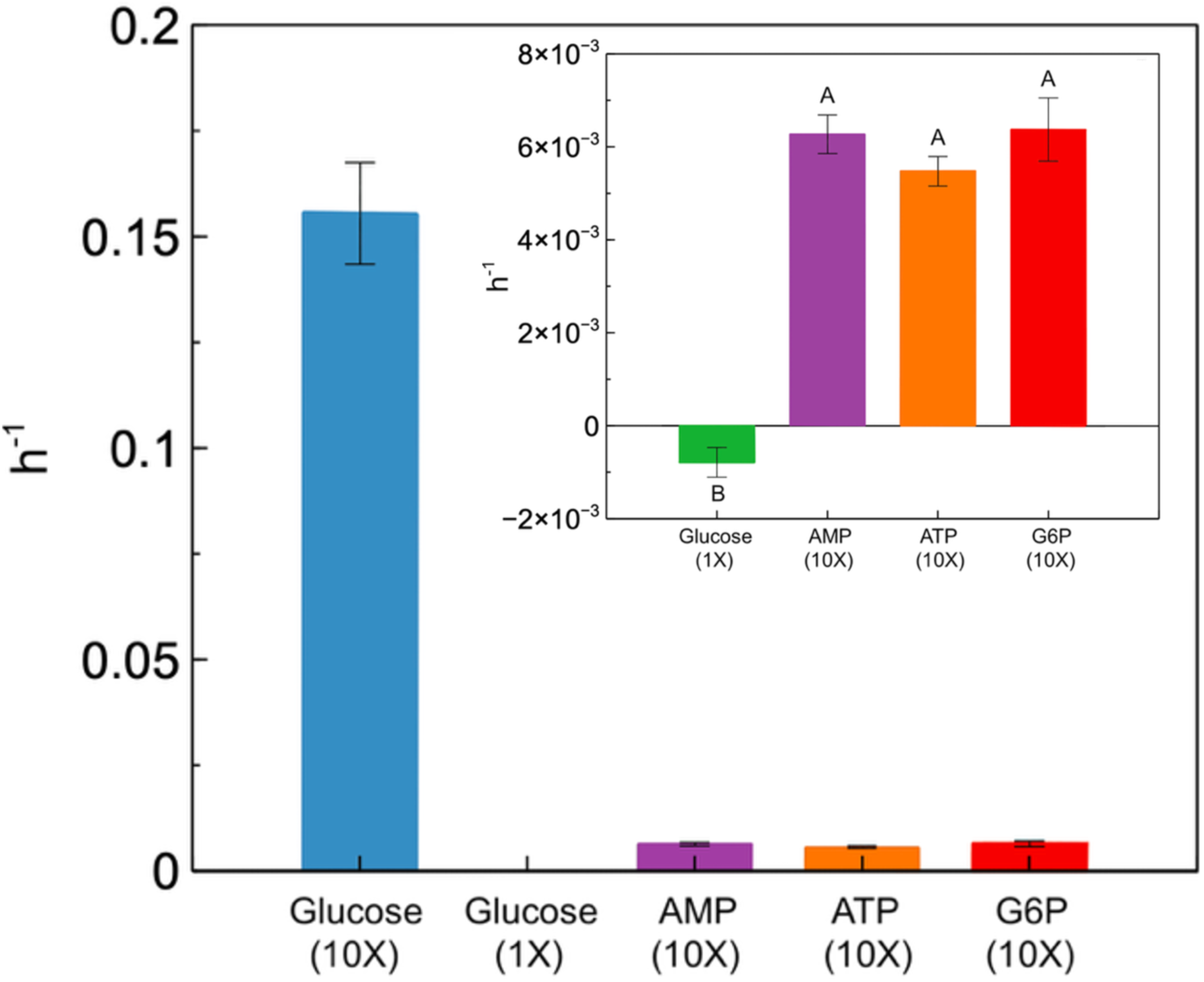
*R. pomeroyi* growth rates. Inset excludes the 10× glucose group for clarity. Groups lacking a shared letter are significantly different (p < 0.05) (Tukey HSD). Error bars represent the standard deviation of the mean for biological triplicates.

Cell-normalized APA was highest in the 1× glucose cultures (6.89 fmol cell^-1^ hr^-1^), followed by G6P (4.79 × 10⁻^1^ fmol cell^-1^ hr^-1^), 10× glucose (9.49 × 10⁻^2^ fmol cell^-1^ hr^-1^), AMP (3.65 × 10⁻^2^ fmol cell^-1^ hr^-1^), and ATP (1.31 × 10⁻^2^ fmol cell^-1^ hr^-1^) (Fig. 8). Final concentrations of SRP were 122 ± 6 µM (10× glucose), 236 ± 4 µM (1× glucose), 293 ± 7 µM (AMP), 289 ± 15 µM (G6P) and 253 ± 17 µM (ATP).

**Figure 8:**
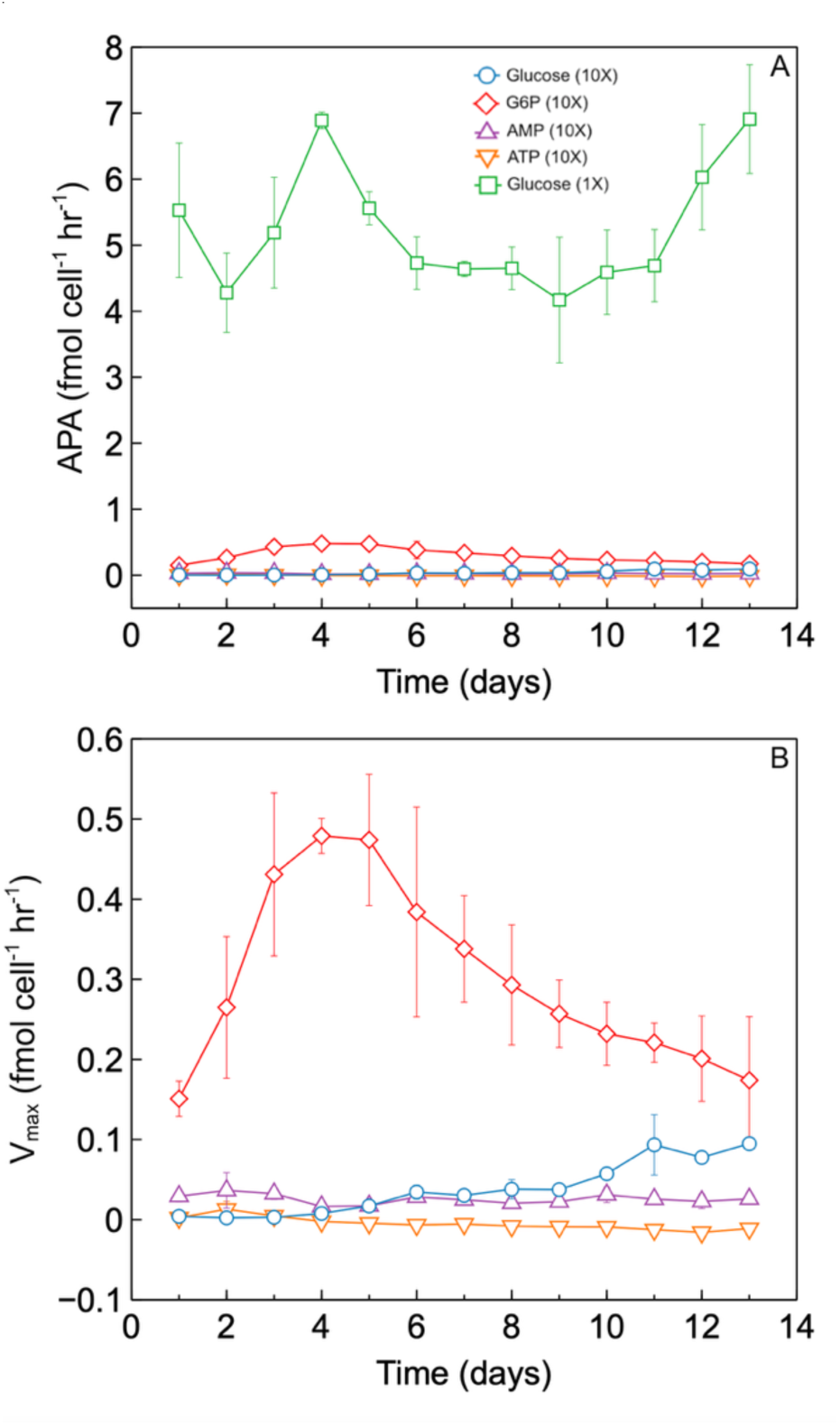
*R. pomeroyi* APA. **(A)** All treatments. **(B)** With the 1× glucose group excluded for clarity. Error bars represent the standard deviation of the mean for biological triplicates. Interpolation lines indicate linear interpolation.

## Discussion

The overall goal of this study was to understand the role of APs in aquatic systems with relatively high phosphate, such as the coastal waters surrounding the Scripps Pier. This study builds on previous work regarding phosphorus hydrolase activity at this location (Ammerman and Azam, 1991). The Scripps Pier environment can be characterized as phosphate-replete over our one-year time series based on the molar DIN:SRP ratios, which were typically below the Redfield value of 16. In addition, DOP represented 56.5 ± 15.4% of total dissolved P throughout the study period, which is less than typically observed in phosphate-stressed conditions (Duhamel et al. 2021). Consistent with the expected seasonality of the system, inorganic nutrients tended to be highest in the winter and lowest in the spring/summer, when phytoplankton biomass, as indicated by chlorophyll, was low and high, respectively.

The observed range of APA (V_max_) values during the year-long period varied from 0.004 to 2.82 nmol P µg Chl-a⁻¹ h⁻¹ (Fig. 4B), which is consistent with findings from other studies in similar regions. A study conducted in the Celtic Sea (coastal northeastern Atlantic) reported APA values ranging from 0.14 to 23.6 nmol P µg Chl-a⁻¹ h⁻¹ (Davis and Mahaffey, 2017), while a study off the coast of Oregon reported values between 0.01 and 0.148 nmol P µg Chl-a⁻¹ h⁻¹ (Ruttenberg and Dyhrman, 2012). Similarly, the K_m_ values observed here (35 to 2615 nM), fall within the range reported for the Celtic Sea surface mixed layer (34 to 3143 nM) (Davis and Mahaffey, 2017) (Fig. 4D). SRP concentrations during the Scripps Pier year-long study ranged from 42 to 702 nM, which is comparable to the Celtic Sea study (30 to 770 nM) and the Oregon coast study (20 to 850 nM) (Davis and Mahaffey, 2017; Dyhrman and Ruttenberg, 2006) (Fig. 3A). Although phosphate is thought to regulate APA (Dyhrman and Ruttenberg, 2006), no clear relationships between SRP and V_max_ or K_m_ were seen at the Scripps Pier (Supplementary Fig. 5).

In the time series, DOP concentrations ranged from ∼40 nM to a peak of 600 nM, which is consistent with other regions (Lin et al. 2012; Liang et al. 2023; Nausch et al. 2018).

Furthermore, DOP concentrations at the Scripps Pier were frequently higher than K_m_ values, suggesting that the substrate-saturated V_max_ values reported here may realistically approximate *in situ* APA, in at least a portion of the dataset. However, we observed no relationships between DOP and V_max_ or K_m_ at the Scripps Pier (Supplementary Fig. 5). Biomass may regulate APA in marine systems and help explain some of the variability observed during the sampling period, as APA and Chl-*a* have shown to exhibit a linear relationship at phosphate concentrations of 0.2–1.0 µM (Nausch, 1998). However, chlorophyll only explained 5% of the variability in APA (Supplementary Fig. 3), and APA remained dynamic even after normalization to Chl-*a* (Fig. 4B).

It is commonly known that DOP can serve as a phosphorus source via the release of phosphate from DOP by AP enzymes. By removing phosphate from the organic molecules, APs may also increase the bioavailability of organic carbon (Karl, 2014). Because APs have been hypothesized to play a role in organic carbon acquisition by heterotrophs (Hoppe and Ullrich, 1999), we tracked DOC concentrations at the Scripps Pier. DOC concentrations were relatively stable over the time series, with a few exceptions, and were consistent with other coastal systems (Archer et al. 1997; De Vittor et al. 2008; Lønborg et al. 2024). Yet we did not observe any relationships between DOC and V_max_ or K_m_ during the one-year time series (Supplementary Fig. 5). If APs are involved in organic carbon acquisition, we hypothesized that APA could be related to the phosphorus content of dissolved organic matter (DOM). The typical DOC:DOP ratio in this study was 278 ± 151. The elevated C:P ratio in the DOM pool (compared to the Redfield ratio of 106:16:1) is well known, with bulk DOC:DOP ratios ranging from 251 (Equatorial Pacific) to 638 (North Atlantic Subtropical Gyre) in the surface ocean (Liang et al. 2023). Given the stability of DOC concentrations throughout the time series, variations in DOP concentrations likely drove changes in DOC:DOP stoichiometry, but we observed no relationships among DOC:DOP and V_max_ or K_m_. DOP is known to be more labile than both DOC and dissolved organic nitrogen (DON), in addition to being remineralized twice as fast as either (Letscher and Moore, 2015; Lønborg & Álvarez-Salgado, 2012). Indeed, some phytoplankton in this region utilize DOP with diverse hydrolase enzymes, despite relatively phosphate-rich conditions (Steck et al. 2025). Both autotrophic and heterotrophic consumption of surface DOP may help explain observed changes seen in DOC:DOP over the sampling period (Liang et al. 2023).

Despite the waters off the Scripps Pier typically being nitrogen-limited and P-replete, APA was detectable for most of the time series, similar to other coastal sites where the APA paradox has been reported. In the absence of any bulk indicators that could account for the APA observed (SRP, DOP, DOC, DOC:DOP, DIN:SRP, chlorophyll), we hypothesize that microbial community composition may play a major role. Community composition was highly variable throughout the sampling period and has been divided into five distinct taxonomic modes: high nutrient winter, spring productive, spring bloom, fall bloom, and post-bloom (Adams, 2025).

These modes were characterized by different dominant taxa, reflecting seasonal environmental changes (Adams, 2025). Specifically, the unclassified *Roseobacteraceae* family showed a high prevalence during the fall and some presence in the post-bloom period. A representative of the *Roseobacteraceae*, *R. pomeroyi*, is known to utilize DOP as a sole phosphorus source via Aps (Adams et al. 2022; Sebastian and Ammerman, 2011), but its ability to utilize DOP as a potential carbon source has not been explored to our knowledge.

To investigate the hypothesis that DOP could serve as a sole carbon source for *R. pomeroyi* via the activity of APs, we grew *R. pomeroyi* on three different DOP sources (AMP, ATP, and G6P) and two concentrations of glucose as positive and negative controls. Over the 13-day experiment, *R. pomeroyi* grew on all DOP sources, with G6P supporting the most growth, followed by AMP and ATP (Figs. 5, 6). G6P supported the most growth, likely due to its labile P-ester bond, which is typically hydrolyzed by APs (Srivastava et al. 2021). AMP was found to have the second highest growth among the DOP sources, likely because it also contains a P-ester bond. Although G6P and AMP have the same type of phosphorus bond, their organic moieties differ, which could reflect differences in carbon bioavailability between glucose and adenosine. ATP showed the lowest growth among the DOP sources, perhaps because it contains one P-ester bond and two P-anhydride bonds that must be broken, potentially by different enzymes. The lower growth observed on all DOP sources compared to the equivalent amount of carbon as glucose could be because growth on DOP is more energetically costly (Thomson et al. 2019). Overall, these results confirm that *R. pomeroyi* can acquire carbon from various DOP sources.

The APA analysis of *R. pomeroyi* cultures suggests that AP is involved in some carbon nutritional pathways. The 10-fold depletion of initial carbon concentrations alone resulted in cell-specific APA at least an order of magnitude higher than all other treatments (Fig. 8A), reflecting the upregulation of APA under carbon stress, potentially indicating scavenging for alternate carbon sources (Chan et al. 2012). Consistent with this response, the G6P cultures upregulated APA compared to the C-replete 10× glucose cultures, suggesting that APs are involved in carbon acquisition from G6P. APA eventually began to increase in the 10× glucose cultures, but was always lower than the G6P cultures (Fig. 8B). This could be because there was sufficient carbon present to temporarily suppress APA until glucose became more depleted. If APs were solely involved in phosphate acquisition during phosphate scarcity, then the elevated phosphate concentrations, which we confirmed were still replete by the end of the experiment, would have inhibited APA in these treatments. Since they did not, our results support a carbon nutritional role for APA in these cases. Indeed, our results for *R. pomeroyi* are consistent with the terrestrial microbe *Flavobacterium johnsoniae*, which uses an AP that is not inhibited by phosphate to grow on phosphorylated carbohydrates as sole carbon sources (Lidbury et al. 2022).

However, APA did not appear to be involved in the utilization of carbon from nucleotides, as APA in the ATP and AMP cultures remained similar or lower than the C-replete 10× glucose control (Fig. 8B). The AMP and ATP cultures exhibited the lowest APA levels, with minimal activity throughout the entire growth period (Fig. 8B). Despite this, AMP and ATP cultures showed higher growth than the deplete group, indicating that *R. pomeroyi* can use the carbon from AMP and ATP (Figs. 5–7). One possible explanation is that *R. pomeroyi* acquires carbon from ATP and AMP using a different enzyme, like 5’-nucleotidase (5’-NT). In an experiment confirming that *R. pomeroyi* can grow on AMP, ATP, and other forms of DOP as a sole phosphorus source, proteomic analysis showed the presence of 5’-NT (Adams et al. 2022). Additionally, a study on the diatom *Phaeodactylum tricornutum* showed that ATP supported growth as a sole phosphorus source without detecting APA, despite an increase in phosphate levels. Transcriptomic and RT-qPCR results revealed that the gene for 5’-NT was expressed in these cultures (Zhang et al. 2023). These findings suggest that multiple P hydrolase pathways may be involved in utilizing DOP as carbon sources, and the absence of APA does not necessarily signal a lack of carbon utilization from DOP.

Overall, our results suggest that bacteria contribute to APA in phosphorus-replete environments to access some phosphate-bound forms of DOC. Indeed, some fraction of the DOC pool may not be readily accessible through simpler pathways, leading microbes to use AP as an “alternative” method for carbon acquisition. This research highlights the potential versatility of APs, suggesting they may have broader nutritional roles than widely recognized. The results of this study help address the APA enigma or paradox, which has been observed in various parts of the global oceans but has eluded complete understanding for several decades. Additionally, this study suggests that APs, and potentially other P hydrolase enzymes, may play a significant role in coupling the carbon and phosphorus cycles.

## Acknowledgments

The authors thank Marley Weiss for collaborating on seawater sample collection and Melissa Carter, Kayla Martin, and Elena Beckhaus for coordinating the sampling efforts. We also thank the Southern California Coastal Ocean Observing System (SCCOOS) HABMAP program for providing access to DIN data. This work was supported by the U.S. National Science Foundation under grant 1948042 (J.M.D.) and the Simons Foundation under grant 678537 (J.M.D.).

## Author contributions

E.S. wrote the paper with input from all authors. E.S., K.B.L., S.P., and C.T. collected field samples, analyzed field samples, and conducted data analysis. E.D.I. analyzed field samples. J.C.A. developed analytical methods. E.S. conducted and analyzed laboratory experiments. J.S.B. and J.M.D. acquired funding and coordinated the project. J.M.D. conceptualized the study.

## Data availability

Data from the SCCOOS HABMAP program can be accessed via the ERDDAP online data access form (https://erddap.sccoos.org/erddap/tabledap/HABs-ScrippsPier.html). Data supporting other findings of this study are available through the Biological and Chemical Oceanography Data Management Office (BCO-DMO) under project number 1948042.

## Declarations

The authors have no financial or non-financial interests to disclose. The authors declare no competing interests.

## Notes

### Competing Interest Statement

The authors have declared no competing interest.

