## Supplementary Information for "Alkaline phosphatase activity supports heterotrophic carbon acquisition in a coastal time series site and a representative marine bacterium"

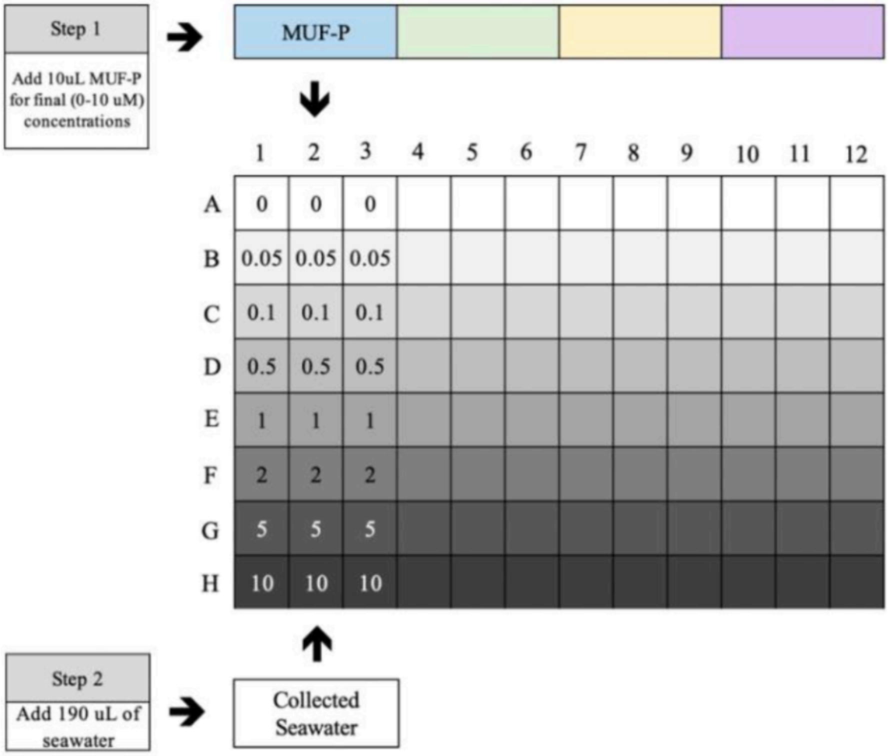


**Supplementary Figure 1:** 96-well plate setup used for determination of alkaline phosphatase activity.

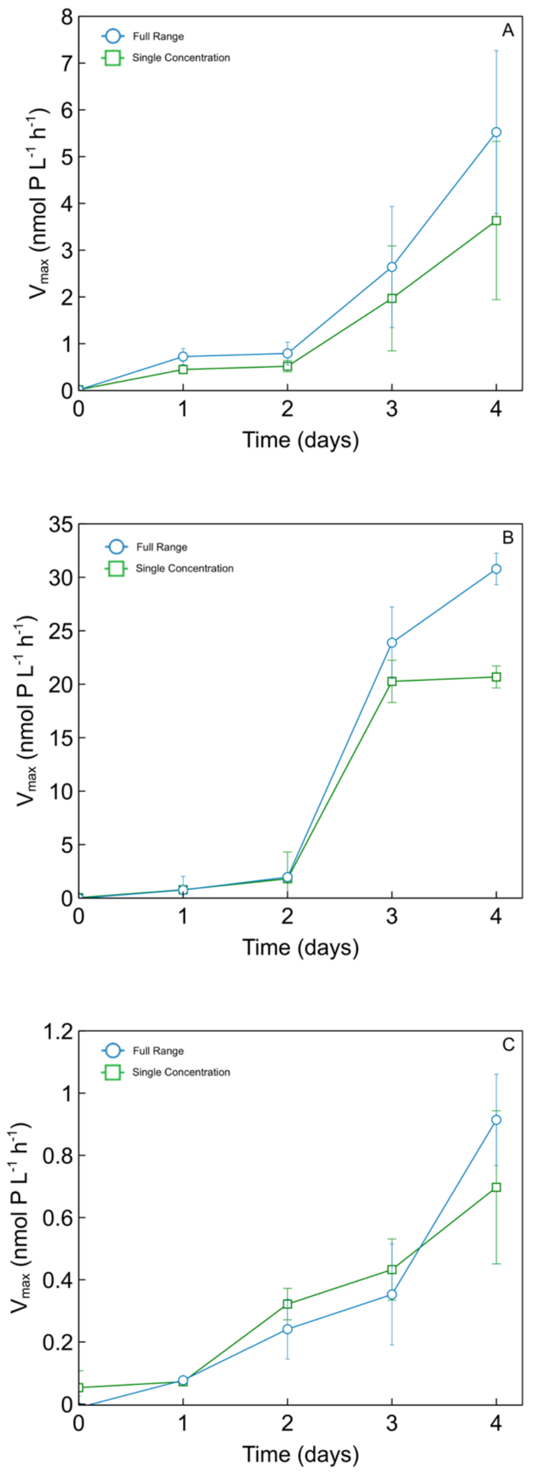


**Supplementary Figure 2:** Alkaline phosphatase activity ($V_{max}$) of *R. pomeroyi* cultures determined using a range of MUF-P concentrations (0-10 μM) vs a single saturating MUF-P concentration (10 μM) in **(A)** Glucose (10×) **(B)** Glucose (1×) **(C)** G6P.

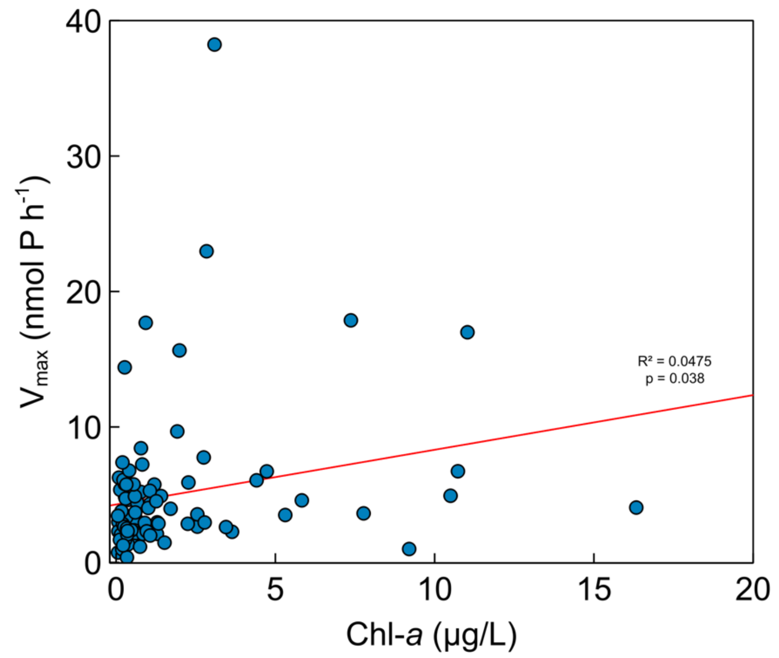


**Supplementary Figure 3:** Volume-normalized $V_{max}$ relative to Chl-*a* at the Scripps Pier.

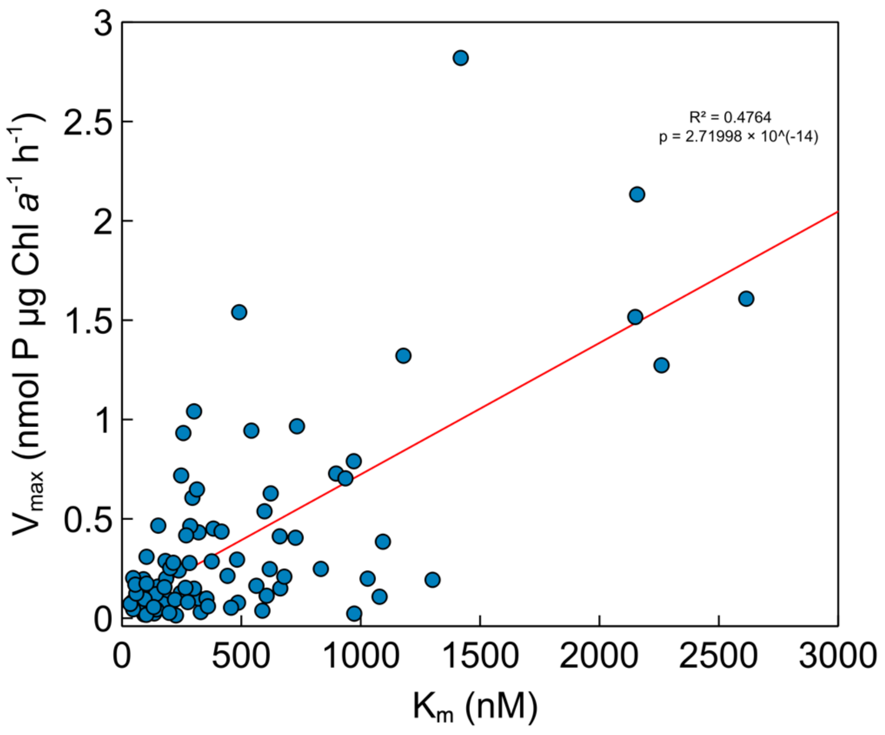


**Supplementary Figure 4:** Chlorophyll-normalized $V_{max}$ relative to $K_m$ at the Scripps Pier.

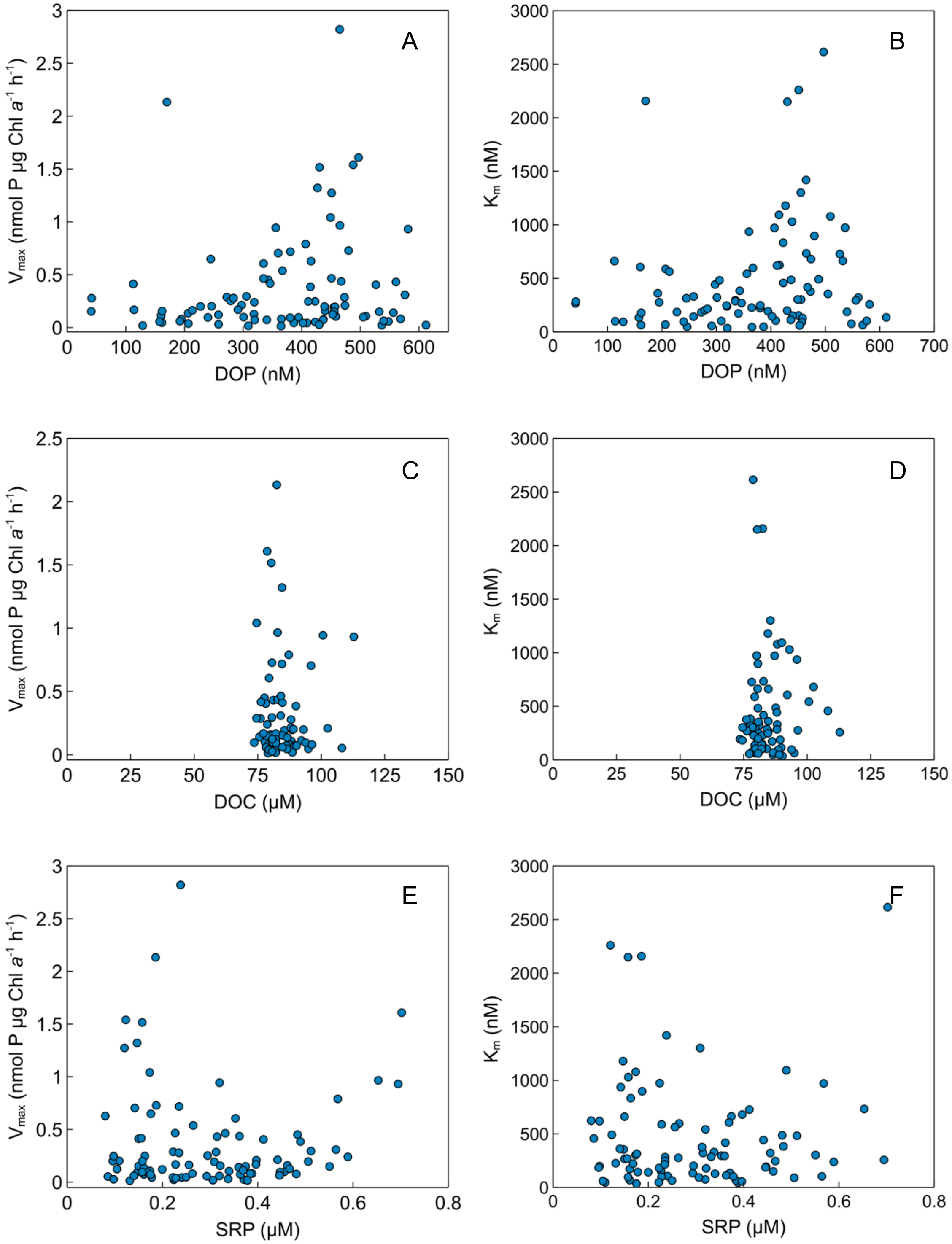


**Supplementary Figure 5:** $V_{max}$, $K_m$, DOP, DOC, and SRP at the Scripps Pier.
